# The conserved NUS-1/DHDDS complex links N-glycosylation to lipid and lysosomal homeostasis in *C. elegans*

**DOI:** 10.64898/2025.12.25.696488

**Authors:** Meixi Gong, Yilin Du, Wentao Zhao, Lingfan Zhu, Changfeng Jia, Yuxiao Xie, Chen Liu, Wenfei Li, Pei Zhao, Zhe Zhang

## Abstract

N-glycosylation, driven by the *cis*-prenyltransferase (*cis*-PT) complex, a NUS1 and DHDDS heterotetramer, is vital for proteome integrity. Despite being linked to severe congenital disorders of glycosylation (CDGs), the systemic role of NUS1/DHDDS remains elusive. Here, we characterize the conserved *C. elegans cis*-PT complex and uncover a critical crosstalk with global lipid metabolism. *cis*-PT deficiency induces catastrophic ER stress, global glycoprotein defects, and developmental failure. Mechanistically, NUS-1 physically interacts with the core oligosaccharyltransferase (OST) complex, directly coupling substrate synthesis to N-glycan transfer. Loss of NUS-1 or DHDDS causes systemic lipopenia, lysosomal dysfunction, and cholesterol sequestration, resulting from impaired N-glycosylation of key regulatory glycoproteins like SCAP. Utilizing CDG-associated CRISPR models, we resolve the molecular basis for *cis*-PT dysfunction, identifying interface disruption and catalytic impairment as distinct disease etiologies. These findings collectively reveal a critical and previously unappreciated crosstalk between protein glycosylation and global lipid metabolism, establishing a robust system to explore CDG pathology and therapeutic strategies.

## Introduction

Maintaining cellular homeostasis is fundamental for the survival and proper functioning of all biological systems, from *E. coli* to humans^1–3^. This intricate balance relies heavily on proteome integrity, a dynamic collection of proteins whose precise folding, modification, and localization are essential for cellular processes^1^. As a common post-translational modification (PTM), N-glycosylation is central to this integrity, significantly influencing glycoprotein structure, stability, trafficking, and function^4,5^. Although N-glycosylation’s fundamental importance in protein function and disease has been extensively studied, its precise roles in broader cellular homeostasis and its interplay with other essential cellular processes remain elusive.

The synthesis of dehydrodolichyl diphosphate (DHDD), the crucial lipid carrier precursor for N-glycosylation, is catalyzed by the *cis*-prenyltransferase (*cis*-PT) complex^6^. This complex is a heterotetramer composed of two essential subunits, nuclear undecaprenyl pyrophosphate synthase 1 (NUS1, also known as NgBR) and dehydrodolichol diphosphate synthetase (DHDDS)^7^. Mutations in NUS1 or DHDDS have been recently linked to severe neurodevelopmental disorders, including epilepsy, intellectual disability, movement disorders, and retinal degeneration, which are classified as congenital disorder of glycosylation (CDG)^7–9^.

While the *cis*-PT complex’s direct role in N-glycosylation is well-established, investigating its full systemic impact is challenging, as complete knockout of its components often causes embryonic lethality in mammalian models, such as mice^9–11^. Consequently, research in other genetically tractable animal models, like *Drosophila* or *zebrafish*, frequently relies on knockdown or haploinsufficiency approaches^9,12,13^, which can limit comprehensive *in vivo* studies of chronic disease progression and developmental-cellular interplay.

Because of its conserved cellular pathways, short lifespan, and well-characterized proteostasis network, *Caenorhabditis elegans* is a prominent model for studying cellular homeostasis and developmental-cellular interplay^9,14,15^. Similar to humans, *C. elegans* shares conserved N-glycosylation pathways, from ER synthesis and quality control to Golgi processing, all essential for development and critical biological functions^16,17^. Recently, *C. elegans* research has substantially advanced our understanding of N-glycosylation’s crucial roles in development, behavior, and neuronal patterning, emphasizing the pathway’s conserved importance across eukaryotes^18–20^. Despite the presence of NUS1 and DHDDS orthologs in *C. elegans*, their specific functions and conserved roles were largely unexplored.

In this study, we characterize the NUS-1/DHDDS complex in *C. elegans*, demonstrating its critical roles in global lipid metabolism and lysosomal homeostasis that extend beyond N-glycosylation. We show that *cis*-PT deficiency induces severe developmental defects, catastrophic ER stress, and widespread glycoprotein defects. Through immunoprecipitation followed by mass spectrometry-based proteomic (IP-MS) analysis, we uncovered a novel physical interaction between NUS-1 and the core oligosaccharyltransferase (OST) complex, directly coupling dolichyl phosphate (Dol-P) synthesis to N-glycan transfer. Furthermore, NUS-1 or DHDDS depletion disrupts cholesterol-lipid metabolism and severely impairs lysosomal function, characterized by elevated lysosomal pH, impaired degradative capacity, and activated autophagy. By creating *C. elegans* CDG disease models via CRISPR-mediated knock-in of human associated mutations (NUS1 R223H and DHDDS R48H)^10,21^, we uncover mechanistic differences underlying *cis*-PT dysfunction. These findings collectively elucidate a critical and previously unappreciated crosstalk between protein glycosylation and global lipid metabolism, establishing a robust *C. elegans* platform for investigating CDG pathogenesis and revealing significant implications for the systemic pathology of glycoprotein-related disorders.

## Results

### NUS-1/DHDDS deficiency drives developmental defects and UPR^ER^ activation

The NUS1/DHDDS complex is responsible for synthesizing long-chain dolichols (DHDD, C_85-100_) by elongating farnesyl diphosphate (FPP, C_15_) with multiple isopentenyl pyrophosphate (IPP, C_5_) units^22^. The resulting DHDD is subsequently processed to form Dol-P, the essential glycosyl carrier required for N-glycosylation (Supplementary Fig. 1a). We identified *F37B12.3* (hereafter *nus-1*), the sole *C. elegans* homolog of mammalian NUS1 (Supplementary Fig. 1b), through a genome-wide RNAi screen for cellular-stress regulators^23^. While the human NUS1 (hNUS1) C-terminal domain possesses prenyltransferase homology to DHDDS and three N-terminal transmembrane (TM) segments (Supplementary Fig. 1b, c)^24,25^, the *C. elegans* NUS-1 contains only one TM segment and lacks an obvious prenyltransferase domain (Supplementary Fig. 1d, e), highlighting a structural divergence in the *cis*-PT.

RNAi knockdown of *nus-1* or *DHDDS* in *C. elegans* resulted in a significantly shorter and paler pleiotropic phenotypes (Fig. 1a, b), similar to knock-out strains (Supplementary Fig. 2a and Supplementary Table 1), suggesting profound defects in fat stores, lipid utilization, or oligosaccharide synthesis^26–28^. Mechanistic investigations revealed that their deficiency led to a pronounced upregulation of *hsp-4*p::GFP (Fig. 1c and Supplementary Fig. 2b), indicating activation of the endoplasmic reticulum unfolded protein response (UPR^ER^)^29^. This activation was specific to the UPR^ER^ and IRE-1 dependent, as loss of IRE-1 prevented the *hsp-4*p::GFP increase, and reporters for *hsp-16.2*p::GFP (cytosolic UPR, UPR^cyto^) and *hsp-60*p::GFP (mitochondrial UPR, UPR^mt^) showed no such upregulation (Supplementary Fig. 2c, d, and Supplementary Table 2)^30,31^. Considering that impaired N-glycosylation promotes misfolded glycoprotein aggregation, these findings link the observed pleiotropic phenotypes directly to ER stress caused by *nus-1/DHDDS* disruption.

**Fig 1.**
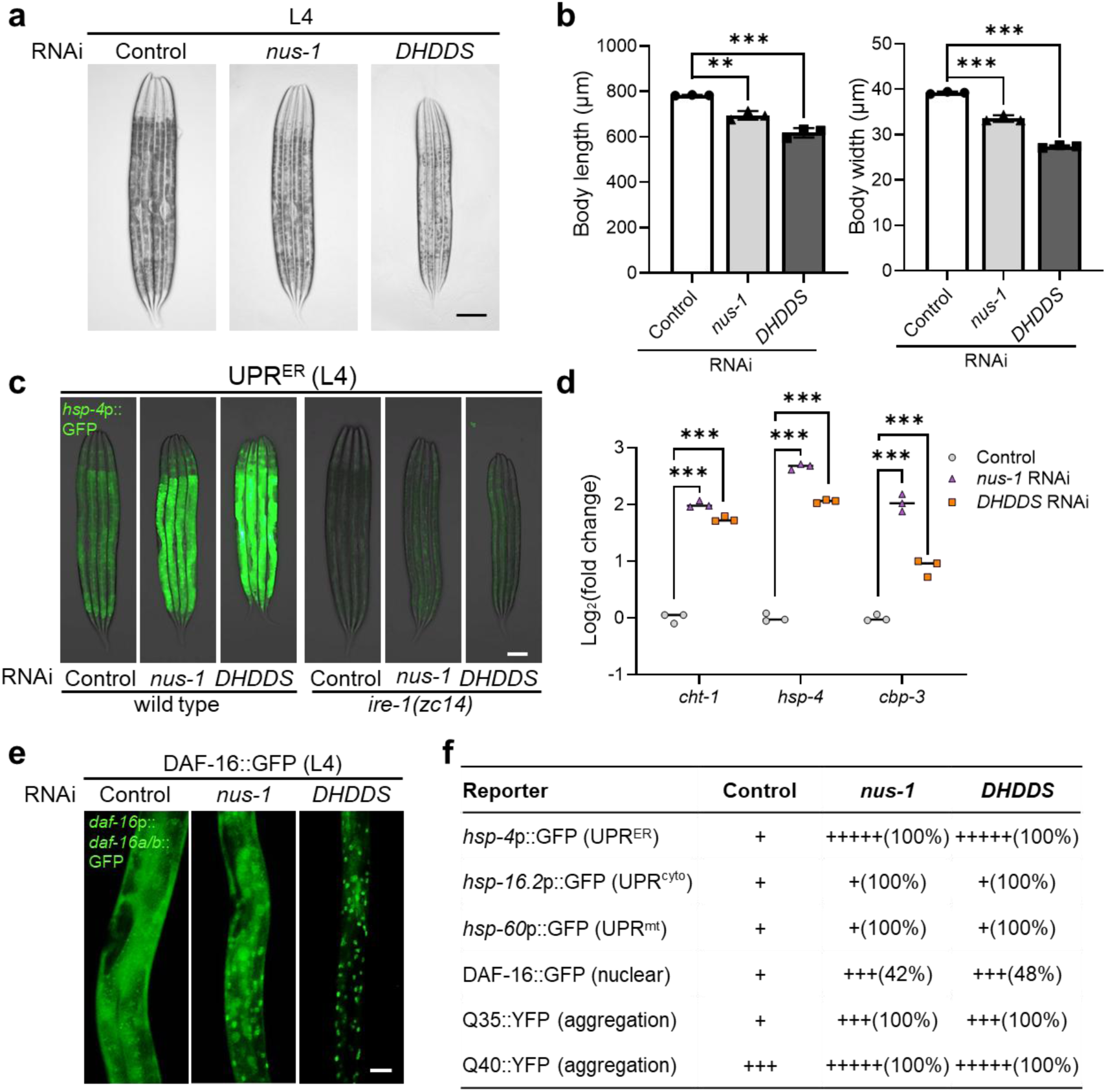
NUS-1/DHDDS deficiency induces developmental defects and activates the UPR^ER^ in *C. elegans*. a, **b** Development defects detection by RNAi against *nus-1* or *DHDDS* at L4 stage with bright-field images (**a**) and quantification (**f**) of animal body lengths and widths (n = 20 for each of the 3 replicates per group, unpaired t-tests: \*\**p* < 0.01, \*\*\**p* < 0.001). Scale bar, 100 μm. **c** Exemplar fluorescence and bright-field images for the UPR^ER^ reporter *hsp-4*p::GFP with *nus-1* and *DHDDS* RNAi in wild-type and *ire-1* mutants. Scale bar, 100 μm. **d** Quantification of *cht-1*, *hsp-4*, and *cbp-3* mRNA fold changes in the indicated RNAi group (n = 3 for each group, unpaired t-tests: \*\*\**p* < 0.001). **e** Fluorescence images of *daf-16*p::*daf-16a/b*::GFP with RNAi against *nus-1* and *DHDDS* at L4 stage. Scale bar, 20 μm. **f** Table listing reporters examined in phenotypic screens for *nus-1* and *DHDDS* RNAi. +, fluorescent reporter levels.

Similar to mammals, the UPR^ER^ in *C. elegans* is initiated by three main pathways, including IRE-1, PEK-1 and ATF-6 (Supplementary Fig. 2e)^32^. Since N-glycosylation defects are known to impact all three UPR branches^33^, we monitored the activity of ATF-6, IRE-1 and PEK-1 using target genes *cht-1*, *hsp-4* and *cbp-3*, respectively (Supplementary Fig. 2e)^34^. Deficiency in either *nus-1* or *DHDDS* caused significant ER stress, activating the UPR through all three sensors (Fig. 1d and Supplementary Fig. 2f). These findings suggest that the role of *cis*-PT complex in regulating N-glycosylation is likely a conserved mechanism in *C. elegans*.

Furthermore, to track this cellular stress, we used the DAF-16::GFP reporter to visualize the activity of a conserved insulin signaling pathway that regulates stress responses in *C. elegans*^35^. Consistent with the UPR^ER^ activation, a deficiency in either *nus-1* or *DHDDS* resulted in a significant nuclear translocation of DAF-16 from the cytoplasm (Fig. 1e and Supplementary Fig. 2g). This breakdown in ER protein quality control was further evidenced by a significant increase in aggregates of aggregation-prone polyglutamine (polyQ) reporters upon *nus-1*/DHDDS knockdown, demonstrating a general loss of cellular proteostasis (Fig. 1f and Supplementary Fig. 2h, i)^36^. Taken together, these findings indicate that NUS-1/DHDDS plays essential roles in protein folding, maintaining the UPR^ER^ and cellular homeostasis in *C. elegans*.

### NUS-1 acts as the ER signal anchor for the *cis*-PT complex

To investigate how NUS-1/DHDDS may regulate ER function and proteostasis, we first examined its expression and localization. Promoter-driven reporters revealed that *nus-1* and *DHDDS* are broadly expressed, with prominent signals in the intestine, hypodermis, and neurons (Fig. 2a, b). Since the *DHDDS* promoter contains six pairs of unique palindromic repeats that prevent conventional in vitro amplification (Supplementary Fig. 3), we generated the DHDDS reporter using an in situ mCherry knock-in (Fig. 2b). In contrast, the *nus-1* reporter was created as an extrachromosomal (Ex) transgene (Fig. 2a and Supplementary Table 1).

**Fig 2.**
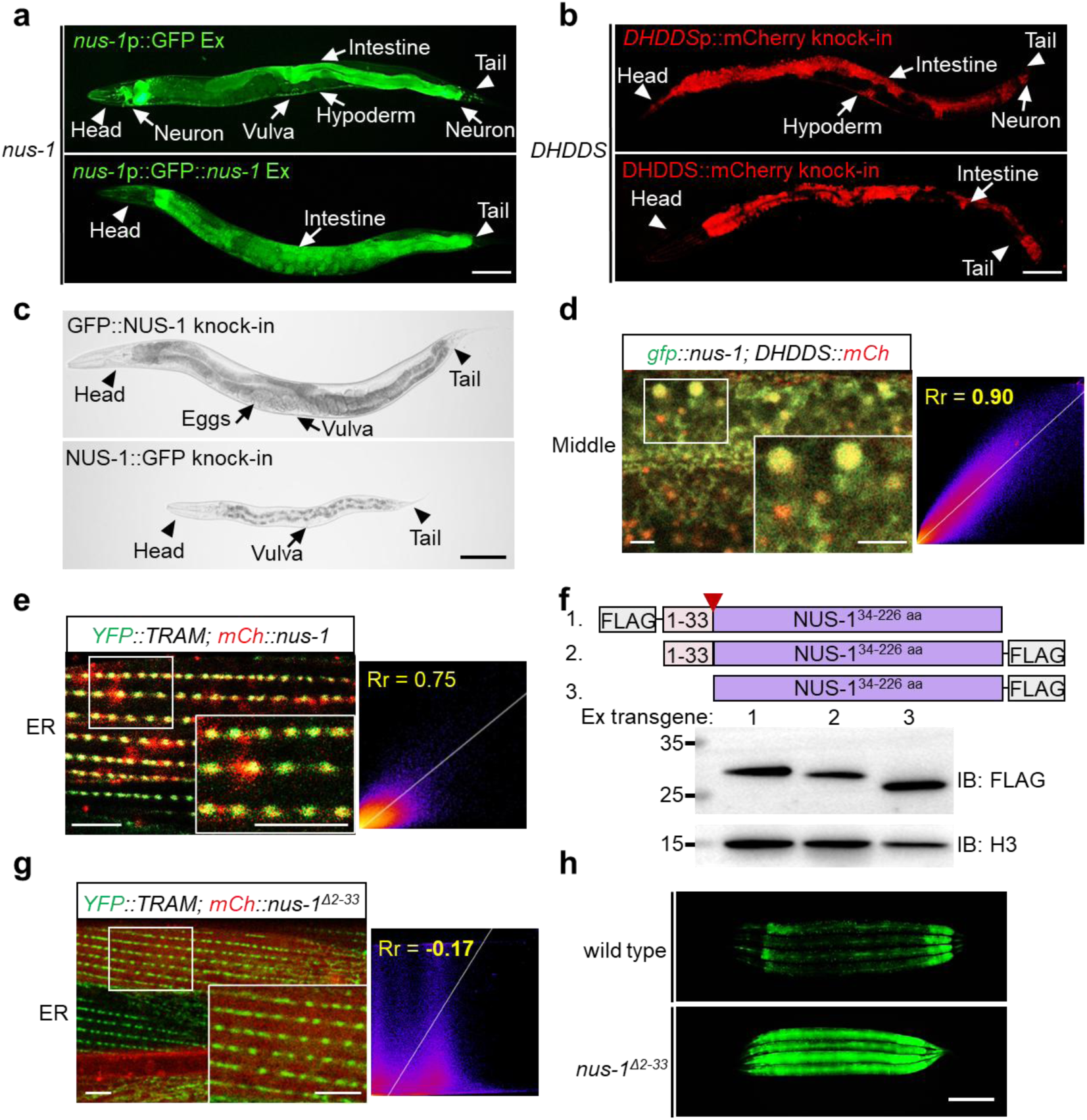
Expression and functional characterization of the *C. elegans* NUS-1/DHDDS complex. a, **b** Expression patterns of NUS-1 (**a**) and DHDDS (**b**) at the D2 adult stage. Panels show transcriptional (upper) and translational (lower) reporters. Ex, extrachromosomal transgene. Scale bars, 100 µm. **c** Representative bright-field images showing the developmental phenotype of animals with in situ GFP knock-ins at the N-terminal (upper) and C-terminal (lower) ends of the *nus-1* gene, respectively. Scale bar, 100 µm. **d, e** Confocal images showing colocalization of GFP::NUS-1 with DHDDS::mCherry (**d**), and mCherry::NUS-1 with the ER marker YFP::TRAM (**e**). Pearson’s correlation coefficient (Rr) indicates linear relationship of pixel intensities (Rr > 0.7 suggests strong colocalization). Scale bar, 5 μm. **f** Western blot analysis of 3xFLAG-tagged NUS-1 constructs. Schematic representation of each fusion protein construct (upper) and Western blot analysis of transgenic strains expressed with expression vectors (lower). The red arrows indicate the predicted signal peptide cleavage site for the first 33 amino acids at the N-terminus. Ex, extrachromosomal transgene. **g** Confocal image showing colocalization of mCherry::NUS-1 with the ER marker YFP::TRAM in the body wall muscle. Rr < 0, negative colocalization. Scale bar, 5 μm. **h** Exemplar fluorescence images for the UPR^ER^ reporter *hsp-4*p::GFP in wild type animals and the NUS-1 signal peptide deletion (NUS-1 ) animals at L4 stage. Scale bar, 100 µm.

When generating in situ GFP-tagged NUS-1 in *C. elegans*, we found that C-terminally tagged animals exhibited growth defects and sterility, while the N-terminally tagged worms were normal (Fig. 2c and Supplementary Fig. 4a, b). This striking phenotypic difference demonstrates that the C-terminus of NUS-1 is functionally critical, mirroring the human NUS1 C-terminal domain essential for DHDDS interaction and *cis*-PT complex activity^37^. Although sequence analysis suggests that *C. elegans* NUS- 1 lacks a distinct prenyltransferase domain (Supplementary Fig. 1b), our results nevertheless indicate a crucial structural and functional role for the C-terminus.

Using functional GFP::NUS-1 and mCherry::DHDDS reporters, validated by their knock-in phenotypes (Fig. 2b, c), we performed in vivo colocalization and subcellular analysis. Our results revealed significant colocalization of these two proteins (Fig. 2d and Supplementary Fig. 4c), indicating they likely form a conserved *cis*-PT complex in *C. elegans*. Further subcellular localization studies showed that mCherry::NUS-1 significantly colocalized with the ER marker YFP::TRAM (Fig. 2e and Supplementary Fig. 4d), but did not substantially colocalize with markers for other major organelles, including the Golgi apparatus, mitochondria, or lysosome (Supplementary Fig. 4e). These results collectively substantiate that the NUS-1/DHDDS complex is a conserved functional unit strictly localized to the ER to maintain cellular homeostasis.

While the N-terminal of human NUS1 serves as a signal anchor (Supplementary Fig. 5a)^38^, the SignalP program predicts that the *C. elegans* ortholog NUS-1 has a cleavable signal peptide (Supplementary Fig. 5b). This raises questions about how it localizes to the ER membrane and anchors the soluble DHDDS protein, which is necessary for dolichol enrichment and N-glycosylation^22,37^. To investigate this, we expressed three 3xFLAG-tagged NUS-1 constructs: one with the tag at the N-terminus, another at the C-terminus of the full-length protein, and a third at the C-terminus of a version lacking the predicted signal peptide (Fig. 2f). Western blot analysis revealed that NUS-1 remains as an uncleaved polypeptide, regardless of the tag’s position (Fig. 2f). Furthermore, deleting the TM domain of NUS-1 in vivo not only disrupted its ER localization (Fig. 2g and Supplementary Fig. 5c) but also caused significant UPR^ER^ activation and severe developmental defects (Fig. 2h and Supplementary Fig. 5d, e). These data unequivocally demonstrate that the non-cleaved TM domain of NUS-1 acts as a signal anchor, essential for ER localization and recruiting the soluble DHDDS subunit to maintain *cis*-PT complex integrity.

### The *cis*-PT complex physically associates with the OST glycosylation machinery

To investigate the physical interactions and homeostatic roles of NUS-1 and DHDDS, we generated transgenic *C. elegans* strains expressing mCherry-tagged variants of both proteins. Affinity purification followed by mass spectrometry (MS) and subsequent co-immunoprecipitation (co-IP) confirmed the physical interaction between NUS-1 and DHDDS in vivo (Fig. 3a-c), consistent with their subcellular colocalization (Fig. 2e). These data provide strong evidence that *C. elegans* NUS-1 and DHDDS form a conserved heterodimeric *cis*-PT complex, analogous to its human counterpart.

**Fig 3.**
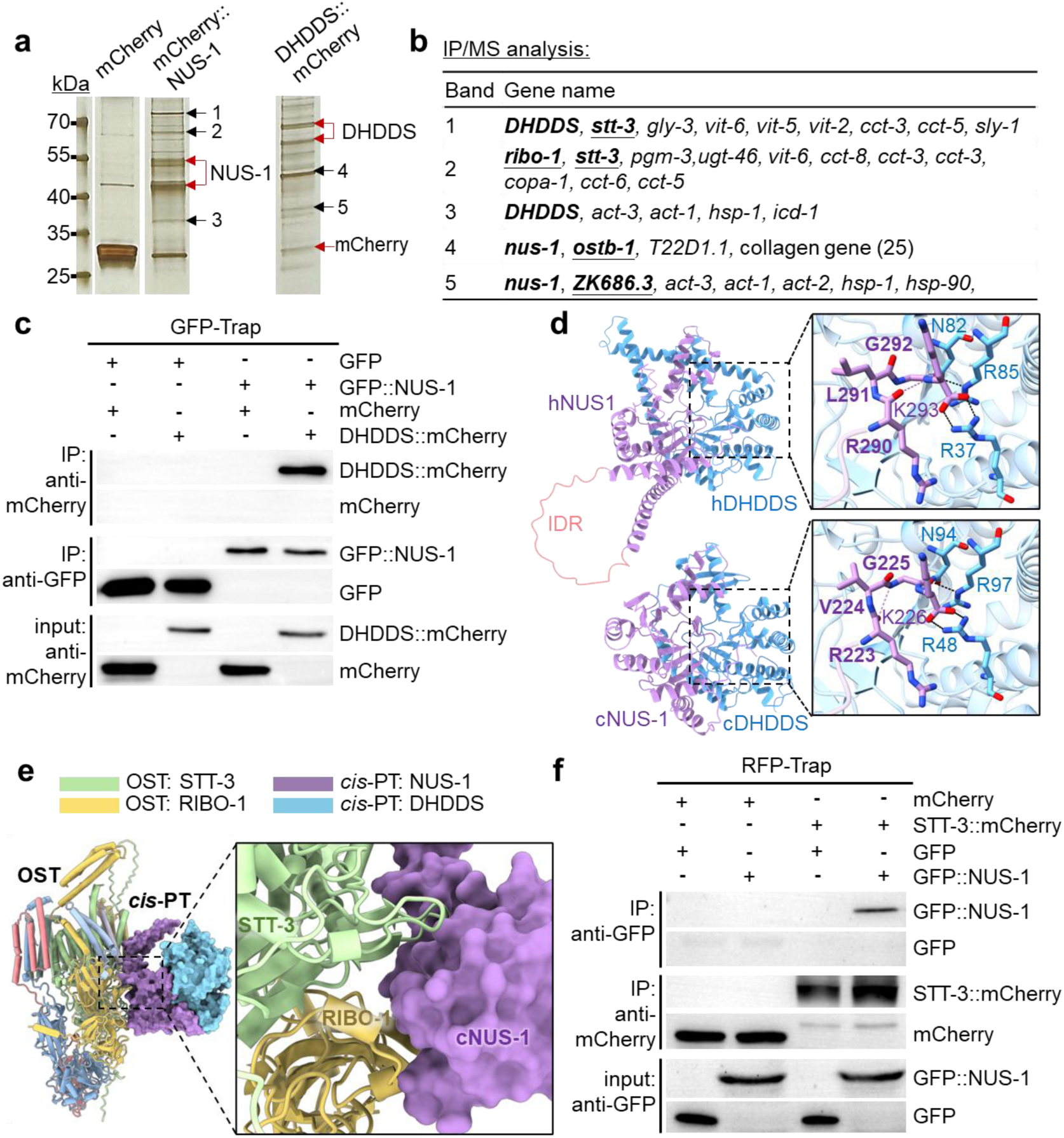
Identification and validation of the NUS-1/DHDDS complex and its interactions. **a** Silver-stained SDS-PAGE showing affinity-purified proteins from total animal lysates expressing mCherry::NUS-1 and DHDDS::mCherry using RFP trap. Red arrows indicate putative NUS-1 and DHDDS bands, and black arrows mark specific bands excised for mass spectrometry (MS) analysis. **b** Table of genes corresponding to proteins identified by MS from unique gel bands (marked as in **a**). *nus-1* and *DHDDS* are shown in bold, and subunits belonging to the OST complex are bolded and underlined. **c** Co-Immunoprecipitation (co-IP) and Western blot confirming the biochemical interaction between N-terminal GFP::NUS-1 and DHDDS::mCherry. Lysates were subjected to IP with GFP-TRAP, followed by immunoblotting with antibodies against GFP and mCherry. **d** Comparative structural analysis of the human (upper) and *C. elegans* (lower) *cis*-PT complexes. NUS1 subunits are violet, and DHDDS subunits are light blue. The intrinsically disordered region (IDR) is shown in salmon red. Insets provide a magnified stick-representation of the conserved catalytic RXG motif, with nitrogen and oxygen atoms in blue and red, respectively. Polar contacts are indicated by dashed lines. **e** Predicted three-dimensional structure model of the *C. elegans cis*-PT/OST supercomplex. Detailed views highlight the interaction interface between the NUS-1 subunit and the OST components STT-3 and RIBO-1. **f** co-IP and Western blot confirming the physical association between GFP::NUS-1 and STT-3::mCherry. Lysates were subjected to IP with RFP-TRAP, followed by immunoblotting with antibodies against GFP and mCherry.

While DHDDS orthologs exhibit high sequence conservation, *C. elegans* NUS-1 shares only 22% identity and 42% similarity with human hNUS1 (Supplementary Fig. 6a, b). Furthermore, NUS-1 lacks canonical prenyltransferase domain found in other species (Supplementary Fig. 1b, c). To reconcile this sequence divergence with functional conservation, we performed the comparative structural modeling of the complex in both *C. elegans* and human (Fig. 3d and Supplementary Movie 1). Although hNUS1 contains a unique N-terminal region composed of two α-helices linked by an intrinsically disordered region (IDR), the rigid domains of both complexes were remarkably similar. We hypothesize that the hNUS1 IDR facilitates the folding of its three N-terminal TM domains, contrasting with the single TM segment identified in NUS-1 (Supplementary Fig. 1d, e). Notably, the catalytic core centered on the RXG motif displayed a nearly identical conformation and residue interactions in both species (Fig. 3d), providing a structural basis for their conserved enzymatic activity.

Analysis of the NUS-1/DHDDS interactome revealed a significant enrichment of glycosylation-related proteins (Fig. 3b and Supplementary Table 3). This cohort included four core subunits of the OST complex (STT-3, RIBO-1, OSTB-1, and ZK868.3)^17,39^, alongside O-glycosylation enzymes (GLY-3, PGM-3) and various glycosyltransferases (UGT-46, T22D1.1). Of particular interest was the OST complex, a conserved enzyme responsible for the final transfer of the glycan to nascent polypeptides (Supplementary Fig. 7a)^40^. Its identification constitutes novel evidence for a direct association between the substrate-producing *cis*-PT complex and the N-glycosylation machinery (Fig. 3b and Supplementary Table 3).

Given this unprecedented finding, we utilized structural modeling to characterize the interface of the putative *cis*-PT/OST supercomplex. Our analysis identified a conserved binding interface within the OST transmembrane region. In *C. elegans*, this interface is primarily mediated by the catalytic subunit STT-3 and the scaffold subunit RIBO-1/RPN1 (Fig. 3e and Supplementary Movie 2), while in humans, it involves STT3B and the accessory subunit MAGT1 (Supplementary Fig. 7b).

Crucially, this interface was predicted to be specific to NUS-1, with the interaction stabilized by a network of hydrogen bonds, whereas no direct binding was predicted for DHDDS (Supplementary Fig. 7c). We functionally validated this model through independent co-IP experiments, which confirmed the physical association between NUS-1 and the catalytic subunit STT-3 (Fig. 3f and Supplementary Fig. 7e). These results suggest that the association of DHDDS with other OST components, such as OSTB-1/DDOST and ZK868.3/MAGT1, is likely mediated indirectly through its partnership with NUS-1. This physical link provides the first evidence for a collaborative functional unit formed in vivo by these sequential enzymes, supporting a model of substrate channeling during N-glycosylation^41^.

### The NUS-1/DHDDS complex is an essential regulator of glycoprotein homeostasis

The interaction of NUS-1/DHDDS with the OST complex prompted us to investigate how N-glycosylation defects impact glycoprotein function and organismal development. We assessed its impact on lysosomal integrity, a process highly dependent on glycoprotein function, using a GFP::GAL3 reporter. GAL3 is a β-galactoside-binding protein that aggregates when it binds to the luminal glycoproteins exposed by lysosomal membrane permeabilization (LMP)^42^. Knocking down either *nus-1* or *DHDDS* resulted in a significant increase in punctate aggregates in the muscle, hypodermis, and intestine (Fig. 4a, b and Supplementary Fig. 8a-c), which directly indicates that NUS-1/DHDDS deficiency disrupts lysosomal membrane integrity.

**Fig 4.**
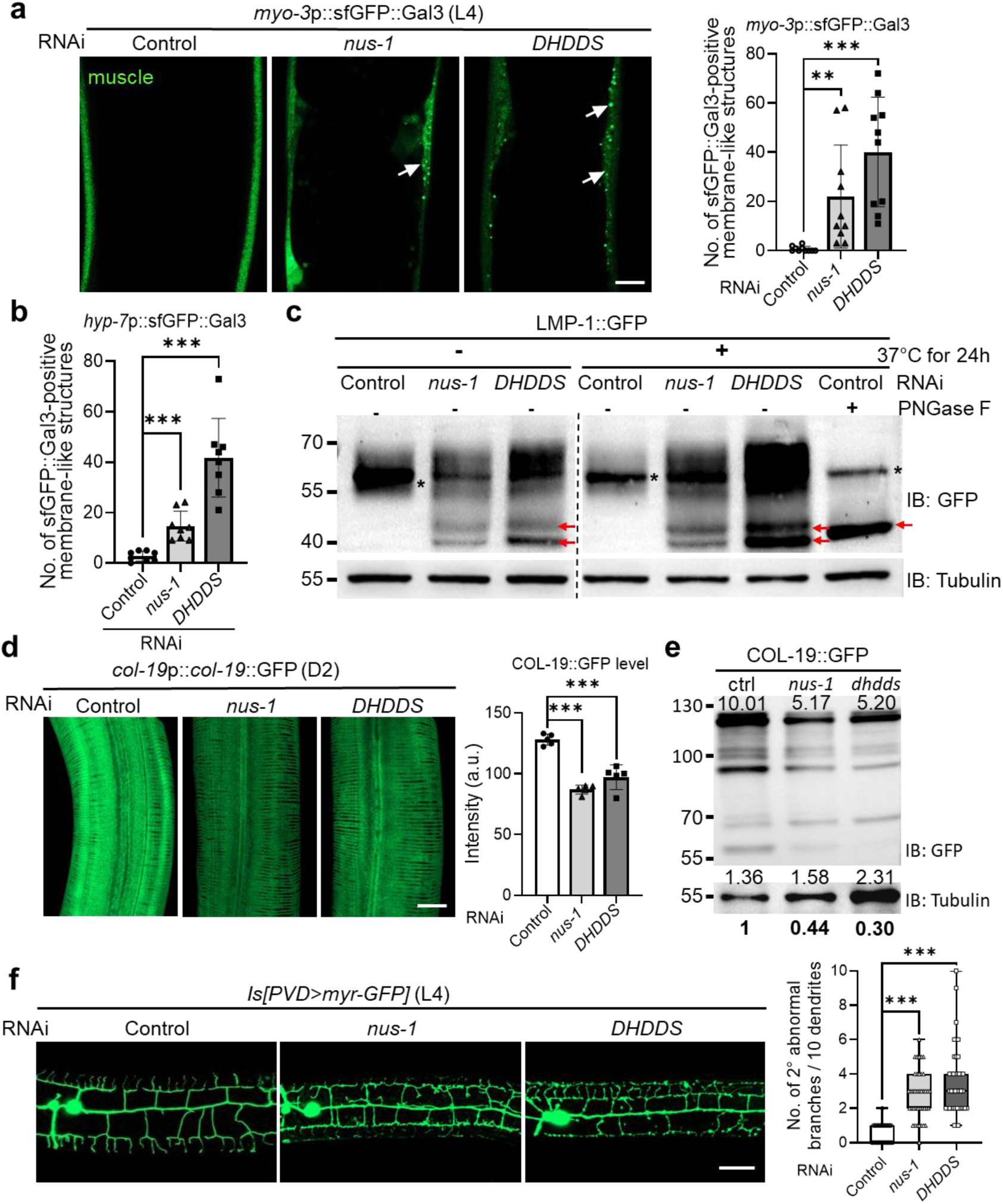
The NUS-1/DHDDS Complex is a conserved regulator of glycoprotein homeostasis in *C. elegans*. **a** Confocal images and quantification of sfGFP::Gal3 aggregates in muscle with RNAi against *nus-1* or *DHDDS* at L4 stage (n = 10 for each group, unpaired t-tests: \*\**p* < 0.01, \*\*\**p* < 0.001). White arrows indicate the sfGFP::Gal3-positive membrane-like structures. Scale bar, 10 μm. **b** Quantification of sfGFP::Gal3-positive aggregates in the hypodermis (*hyp-7p*) following *nus-1* or *DHDDS* RNAi at L4 stage (n = 8 for each group, unpaired t-tests: \*\*\**p* < 0.001). **c** Representative Western blot analysis of lysates from transgenic animals expressing LMP-1::GFP. *C. elegans* were subjected to RNAi against Control, *nus-1*, and *DHDDS*. The samples were either mock-treated or treated with PNGase F, an enzyme that cleaves N-linked oligosaccharides. The higher molecular weight band (asterisk) corresponds to the glycosylated form of LMP-1::GFP. A lower molecular weight band (red arrow) accumulates after treatment, indicating deglycosylation. IB, immunoblotting. **d** Confocal images and quantification of COL-19::GFP with RNAi against *nus-1* and *DHDDS* at D2 stage (n = 5 for each group, unpaired t-tests: \*\*\**p* < 0.001). Scale bar, 20 μm. a.u., arbitrary units. **e** Representative Western blot showing COL-19::GFP protein levels in total animal lysates. Numbers above the blot represent the raw densitometric quantification of each COL-19::GFP and histone H3 band using ImageJ. The relative levels of COL-19::GFP, normalized to histone H3, are shown below the blot in bold. IB, immunoblotting. **f** Loss of *nus-1* or *DHDDS* causes PVD neurodendritic defects. Confocal images of PVD neurons in L4 stage *C. elegans*, visualized by the integrated transgene *ser-2p::myr::GFP* (left). Quantification of abnormal grade 2 PVD neurodendrites per 10 dendrites (right). n = 40 for each group, unpaired t-tests: \*\*\**p* < 0.001. IS, integrated transgene. Scale bar, 20 μm.

To determine if the observed lysosomal membrane disruption was due to unstable glycoproteins, we next examined an LMP-1::GFP reporter strain. The LMP-1 is a lysosome-associated membrane protein (LAMP) with five putative glycosylation sites, whose N-glycosylation is essential for its stability^43,44^. Consistent with our hypothesis, Western blot analysis of *nus-1* or *DHDDS* knockdown animals removed several high molecular mass bands of the LMP-1::GFP protein that were similar to samples treated with PNGase F^42^, an enzyme that specifically removes N-linked glycans (Fig. 4c and Supplementary Fig. 8d). This finding demonstrates that NUS-1/DHDDS deficiency leads to the instability of glycosylated proteins due to impaired N-glycosylation.

Our IP-MS results identified a strong interaction of DHDDS with collagen proteins (Fig. 3b and Supplementary Table 3), suggesting collagen is a potential glycoprotein substrate and prompting us to examine the effects of *nus-1* and *DHDDS* knockdown on collagen proteins. Given that collagen requires N-glycosylation for thermal and mechanical stability^45^, and that impaired collagen synthesis leads to a short, fragile body similar to the phenotype we observed (Fig. 1a, b), we used COL-19::GFP and COL-101::GFP reporter strains.

We found that *nus-1/DHDDS* deficiency caused a significant reduction in both COL-19::GFP and COL-101::GFP expression (Fig. 4d, e and Supplementary Fig. 8e), and also induced the aggregation and mislocalization of COL-101::GFP (Supplementary Fig. 8f, g). Consistent with our IP-MS results, we also observed a significant accumulation of the yolk protein VIT-2::GFP in the intestine and pseudocoelom of knockdown animals (Supplementary Fig. 8h, i), consistent with its reliance on N-glycosylation for proper folding and transport. These observations demonstrate that NUS-1/DHDDS is essential for maintaining protein homeostasis and transport by facilitating proper N-glycosylation.

Beyond its effects on lysosomal integrity and protein homeostasis, we observed severe developmental defects in the complex dendritic arbor of the PVD neurons. Since N-glycosylation is important for dendrite development^20^, we found that knocking down *nus-1* or *DHDDS* resulted in pronounced branching defects, including disorganized neurite branching and the formation of neuronal knots (Fig. 4f). These broad morphological and protein homeostasis defects underscore the essential role of the *cis*-PT complex in N-glycosylation.

### NUS-1/DHDDS deficiency disrupts lysosomal and autophagic homeostasis

The compromised glycoprotein stability and resulting lysosomal membrane integrity caused by NUS-1/DHDDS deficiency prompted a deeper investigation into the resulting lysosomal phenotypes. We first examined the PGP-2::GFP reporter strain, which labels the membrane of lysosome-related organelles (LROs) in *C. elegans*^46^. Consistent with disrupted lysosomal membrane integrity, *nus-1* or *DHDDS* knockdown led to severely swollen LROs and a significant reduction in their number (Fig. 5a, b and Supplementary Fig. 9a). We further corroborated this finding with the lysosomal reporter NUC-1::mCherry^47^, which revealed a profound loss of vesicular integrity and widespread cytoplasmic diffusion of its signal (Fig. 5c and Supplementary Fig. 9b), confirming protein leakage. Concurrently, the significant reduction in the number of NUC-1-positive vesicles (Supplementary Fig. 9c) served as a clear sign of organelle loss or degradation.

**Fig 5.**
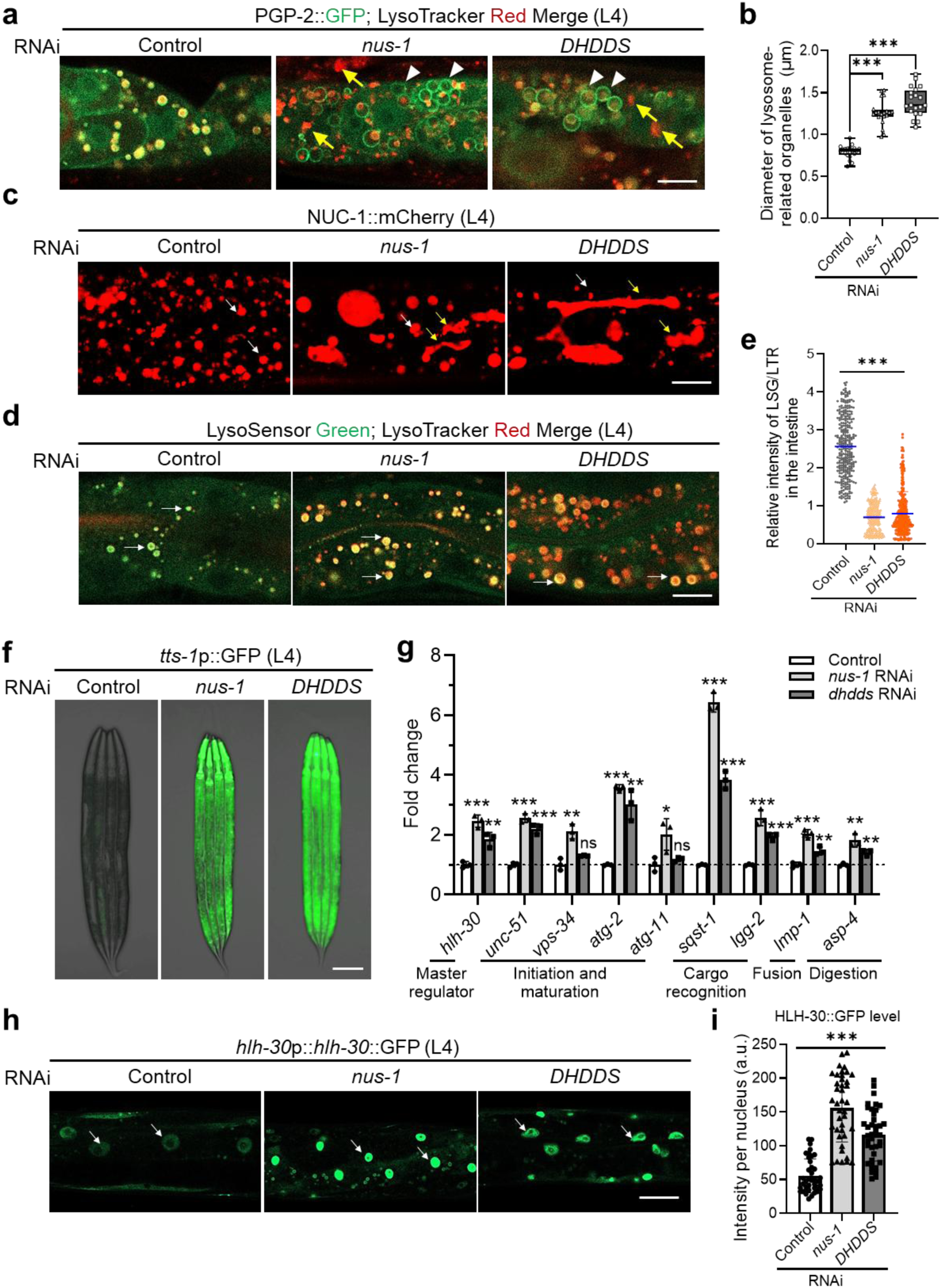
Defects in NUS-1 and DHDDS lead to lysosomal dysfunction and altered autophagy in *C. elegans*. a, **b** PGP-2::GFP and LysoTracker Red co-staining reveals enlarged lysosomes in *nus-1*/*DHDDS* RNAi animals at L4 stage. Representative confocal images (**a**) show expanded PGP-2::GFP-labeled organelles (white arrowheads) and extralysosomal acidic regions (yellow arrows). Quantification of the diameter (**b**) confirms significant enlargement. Scale bar, 10 μm. n = 20 for each group, unpaired t-tests: \*\*\**p* < 0.001. **c** Confocal images show NUC-1::mCherry localization in L4-stage animals with RNAi treatment. White arrows, intact lysosomes. Yellow arrows, signs of lysosomal leakage. Scale bar, 10 μm. **d, e** Representative confocal images (**d**) and quantification of the relative intensity ratio (**e**) of LSG DND-189/LTR DND-99 co-stained intestines in L4-stage animals following RNAi knockdown. White arrowheads indicate vesicular lysosomes stained by both LSG and LTR. Scale bar, 10 μm. Relative intensity of LSG/LTR was quantified (n = 25 per group, 12 lysosomes were quantified per animal, unpaired t-tests: \*\*\**p* < 0.001). **f** Exemplar epifluorescence images of *tts-1*p::GFP with RNAi against *nus-1* and *DHDDS* at L4 stage. Scale bar, 100 μm. **g** Transcriptional analysis of key autophagy genes in the indicated RNAi groups. Gene expression was quantified by RT-qPCR (n = 3 per group, unpaired t-tests: \**p* < 0.05, \*\**p* < 0.01, \*\*\**p* < 0.001, ns, no significant difference). **h, i** Confocal images (**h**) and quantification of the nuclear intensity (**i**) of HLH-30::GFP in L4-stage with RNAi against *nus-1* and DHDDS. White arrows indicate the nuclear localization of HLH-30::GFP. Scale bar, 10 μm. n = 8 animals per group, with at least 4 nuclei quantified per animal, unpaired t-tests: \*\*\**p* < 0.001. a.u., arbitrary units.

To directly assess lysosomal function, we next measured both acidification and degradative capacity. Using the heat-shock inducible pH-sensitive reporter NUC-1::pHTomato, we observed significantly higher fluorescence intensity in *nus-1* or *DHDDS* knockdown animals (Supplementary Fig. 9d, e), directly indicating an elevation in lysosomal pH^47,48^. This was further confirmed by a ratiometric analysis with LysoTracker Red DND-99 (LTR) and the acidotropic dyes LysoSensor Green DND-189 (LSG)^49^. The decreased LSG/LTR intensity ratio, a direct measure of lysosomal acidity, provided compelling evidence that lysosomal pH had risen due to the leakage of acidic contents (Fig. 5d, e). The change in pH was accompanied by impaired degradative activity^50^, as Western blot analysis of the NUC-1::mCherry reporter showed a significant reduction in the proportion of cleaved mCherry, confirming that the activity of acidic hydrolases was compromised (Supplementary Fig. 9f).

Given that lysosomal dysfunction often triggers a compensatory autophagic response^51^, we next assessed the effect of *nus-1/DHDDS* knockdown on autophagy. Using the autophagy transcriptional reporter *tts-1*p::GFP^52^, we observed a significant activation of this pathway (Fig. 5f, Supplementary Fig. 9g). This was further supported by the notable upregulation of key autophagy-related genes, including the master transcription factor *hlh-30*, and genes involved in autophagosome formation and lysosomal degradation (Fig. 5g, Supplementary Fig. 9h). Consistent with these findings, knocking down *nus-1* or *DHDDS* enhanced the nuclear translocation and expression of the HLH-30::GFP translational reporter (Fig. 5h, i and Supplementary Fig. 9i), providing further evidence of autophagy activation^53^.

To determine if this autophagy activation was a specific consequence of N-glycosylation defects, we performed RNAi knockdown of various glycosylation-related genes identified in our IP-MS interactome analysis. We found that only the depletion of N-glycosylation pathway genes, such as the OST complex, fully recapitulated the autophagy activation, unlike the knockdown of O-glycosylation or other general glycosyltransferases genes (Supplementary Fig. 9j). We then confirmed autophagic flux using the dual-fluorescence reporter mCherry::GFP::LGG-1^54^, which showed a significant shift from yellow to red puncta when the pH-sensitive GFP signal was quenched upon autophagosome-lysosome fusion (Supplementary Fig. 9k, l). Collectively, our results demonstrate that NUS-1/DHDDS deficiency severely impairs lysosomal function, which in turn activates the N-glycosylation-dependent autophagic response, highlighting a critical breakdown in cellular homeostasis.

### Disruption of N-glycosylation impairs lipid and cholesterol homeostasis

Beyond the severe developmental and activation of UPR^ER^, the characteristic short and pale body morphology of NUS-1/DHDDS deficiency strongly suggested an underlying defect in lipid metabolism^27^. We first investigated neutral lipid storage using the lipophilic dye Oil Red O^55^, and observed a significant, global reduction in total lipid content across *cis*-PT deficient animals (Fig. 6a). This severe deficiency of lipids (lipopenia) was corroborated by a dramatic reduction in both the size and number of neutral lipid droplets, the primary storage sites in *C. elegans*, as visualized by the DHS-3::GFP reporter (Fig. 6b, c and Supplementary Fig. 10a), clearly establishing a fundamental defect in lipid utilization or synthesis^56^.

**Fig 6.**
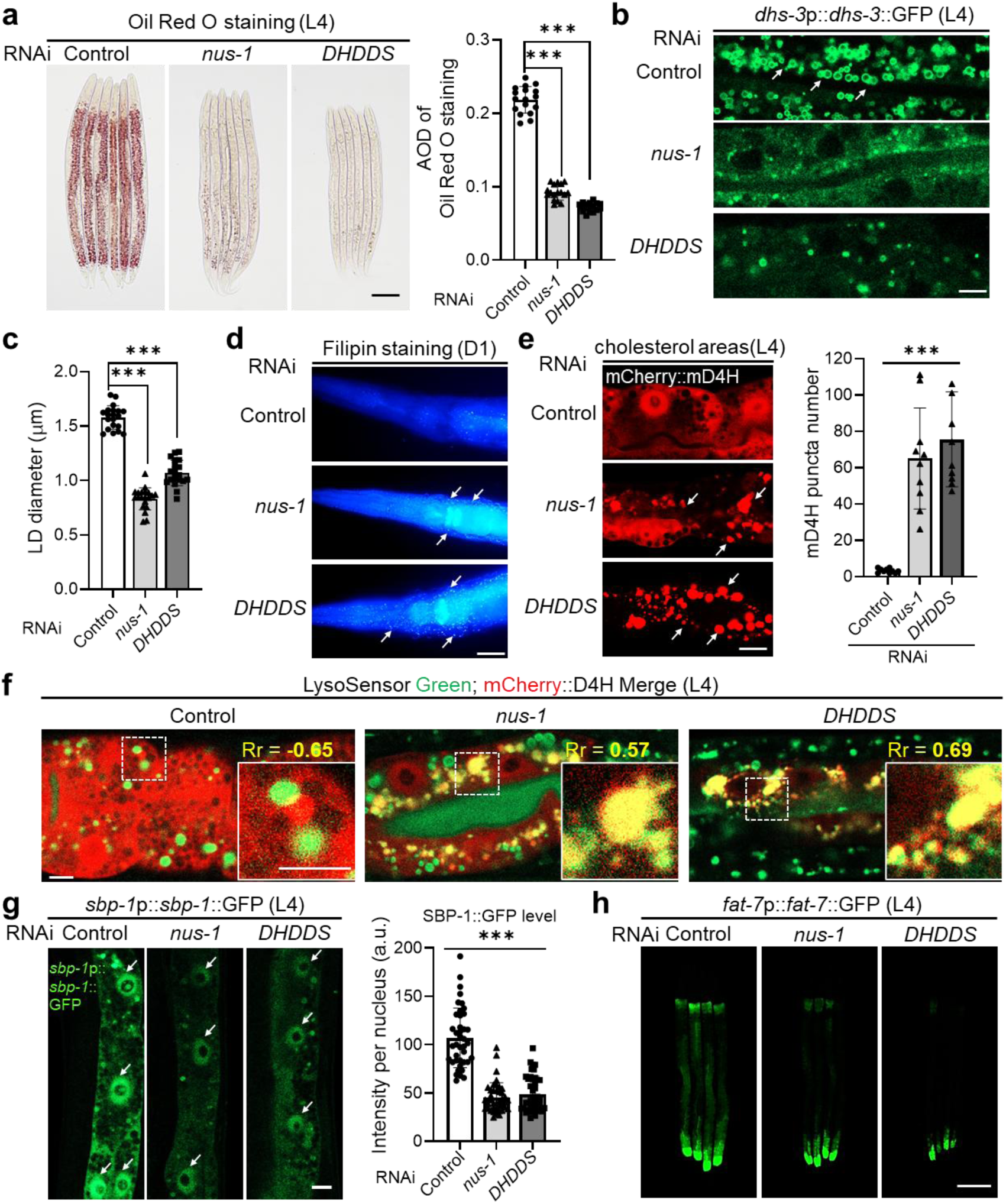
NUS-1/DHDDS deficiency disrupts lipid metabolism and cholesterol homeostasis. **a** Bright-field images of Oil Red O staining of animals with *nus-1*/*DHDDS* RNAi at the L4 stage. Average optical density (AOD) was quantified for each group (n = 20 per group; unpaired t-tests: \*\*\**p* < 0.001). Scale bar, 100 μm. **b, c** Confocal images of DHS-3::GFP-labeled lipid droplets (**b**) and quantification of their diameter (**c**) in animals fed with *nus-1/DHDDS* RNAi. White arrowheads indicate lipid droplets. Scale bar, 10 μm. n = 20 per group. unpaired t-tests: \*\*\**p* < 0.001. **d** Representative fluorescence images of Filipin staining in *nus-1*/*DHDDS* RNAi animals at D1 adult stage. White arrows indicate the accumulation of un-esterified cholesterol. Scale bar, 20 μm. **e** Confocal images of mCherry::D4H-labeled cholesterol (left) and quantification of its aggregation (right) in *nus-1/DHDDS* RNAi animals. White arrowheads indicate cytoplasmic puncta of cholesterol accumulation. Scale bar, 10 μm. n = 10 per group. unpaired t-tests: \*\*\**p* < 0.001. **f** Confocal images showing colocalization of LSG DND-189 with mCherry::D4H. Pearson’s correlation coefficient (Rr) indicates linear relationship of pixel intensities (Rr < 0 suggests negative colocalization, Rr > 0.5 suggests good colocalization). Scale bar, 5 μm. **g** Confocal images of SBP-1::GFP nuclear translocation (left) and quantification of its nuclear fluorescence (right) in L4-stage animals with *nus-1/DHDDS* RNAi. Scale bar, 10 μm. a.u., arbitrary units. n = 8 per group, with 5 nuclei quantified per animal. unpaired t-tests: \*\*\**p* < 0.001. **h** Exemplar epifluorescence images of FAT-7::GFP with RNAi against *nus-1*/*DHDDS* at L4 stage. Scale bar, 100 μm.

Prompted by the severe lipid metabolism defect, we next investigated its link to cholesterol, a crucial cellular lipid that shares the upstream mevalonate pathway with the *cis*-PT products^57,58^. To precisely determine how *nus-1/DHDDS* deficiency impacts cholesterol content and distribution, we employed two complementary methods: in vitro Filipin staining and in vivo D4H cholesterol binding protein observation. Filipin staining, which specifically marks un-esterified cholesterol in intracellular membranes^59^, revealed a striking accumulation of cholesterol within the cytoplasm of knockdown animals (Fig. 6d and Supplementary Fig. 10b). This abnormal distribution was further confirmed by live imaging using the mCherry::D4H reporter, a sensitive tool for monitoring membrane-associated cholesterol^60^. The reporter signal exhibited distinct, intense cytoplasmic puncta in *cis-PT* knockdown animals (Fig. 6e). Crucially, these mCherry-labeled puncta showed significant colocalization with the acidotropic dye LysoSensor Green, providing compelling evidence that the accumulated cholesterol is sequestered within acidic, lysosome-related organelles (Fig. 6f). These findings suggest that NUS-1/DHDDS deficiency impairs intracellular cholesterol trafficking and distribution, which we propose is a key factor for the severe reduction in neutral lipids.

Since intracellular lipid and cholesterol metabolism are transcriptionally regulated^61^, we next focused on the master transcription factor SBP-1, the homolog of mammalian SREBP (sterol regulatory element-binding protein). Our analysis revealed that *nus-1/DHDDS* deficiency significantly reduced the nuclear localization of SBP-1 (Fig. 6g). This impairment was accompanied by a strong suppression of its downstream target, the Δ9 fatty acid desaturase FAT-7, evidenced by a decrease in fluorescence from the FAT-7::GFP translational reporter (Fig. 6h and Supplementary Fig. 10c). This failure of SBP-1 nuclear translocation and its subsequent downstream transcriptional activity provide a molecular mechanism for the observed lipid metabolism defects.

A striking paradox emerged from our analysis: although *nus-1/DHDDS* knockdown strongly suppressed *fat-7* expression, it significantly upregulated the transcription of its master regulator, *sbp-1* (Supplementary Fig. 10d). This upregulation likely reflects a futile compensatory transcriptional response to severe lipid defects. We propose that SREBP cleavage-activating protein (SCAP), the *C. elegans* homolog SCP-1, an essential glycoprotein for SBP-1’s proteolytic activation and nuclear translocation, is compromised without proper N-glycosylation (Supplementary Fig. 10e)^62^. This uncoupling of *sbp-1* transcription from its protein function provides a direct molecular basis for the observed lipid defects.

To confirm the specificity of this broad phenotypic impact, we performed RNAi knockdown of three classes of glycosylation-related genes identified in our IP-MS analysis (Fig. 3b and Supplementary Table 3). Our results consistently showed that only the depletion of N-glycosylation pathway genes, such as the OST complex, fully recapitulated the multi-faceted NUS-1/DHDDS deficiency phenotypes, including the short/pale body morphology, severe lipid and cholesterol defects via the master transcription factor SBP-1 (Table 1 and Supplementary Fig. 10f-j). This high degree of specificity underscores the central role of N-glycosylation in maintaining cellular homeostasis and provides compelling evidence that the NUS-1/DHDDS complex regulates lipid metabolism by ensuring the proper function of essential glycoproteins like SCAP, which is critical for SBP-1 activity.

**Table 1.** Phenotypic consequences of glycosylation gene depletion on lipid and cellular homeostasis. The degree of phenotypic change is indicated by symbols: + denotes increasing activation (HLH-30::GFP, SBP-1::GFP, FAT-7::GFP induction); - denotes decreasing levels (Oil Red O, DHS-3::GFP size reduction) or reporter reduction (FAT-7::GFP); N.E. indicates no observable effect.

| Gene | Function | HLH-30::GFP | DHS-3::GFP | Oil Red O | SBP-1::GFP | FAT-7::GFP |
| --- | --- | --- | --- | --- | --- | --- |
| *nus-1* | *cis*-PT, N-glycosylation | +++ | --- | --- | --- | --- |
| *DHDDS* | *cis*-PT, N-glycosylation | +++ | --- | --- | --- | --- |
| *stt-3* | OST, N-glycosylation | +++ | --- | --- | --- | --- |
| *ribo-1* | OST, N-glycosylation | +++ | --- | --- | --- | --- |
| *ostb-1* | OST, N-glycosylation | +++ | --- | --- | --- | --- |
| *ZK868.3* | OST, N-glycosylation | +++ | --- | --- | --- | --- |
| *gly-3* | O-glycosylation | N.E. | N.E. | N.E. | N.E. | N.E. |
| *pgm-3* | O- glycosylation | N.E. | ++ | ++ | -- | - |
| *ugt-46* | glycosyltransferase | N.E. | N.E. | N.E. | N.E. | N.E. |
| *T22D1.1* | glycosyltransferase | + | + | + | - | N.E. |

### Structural and phenotypic characterization of pathogenic *cis*-PT mutations in the *C. elegans* model

Having demonstrated the essential function of the *cis*-PT complex in glycosylation and lipid homeostasis in *C. elegans*, we next sought to establish in vivo NUS-1/DHDDS-associated disease models. Human mutations in NUS1 and DHDDS are linked to severe congenital disorder of glycosylation (CDG) phenotypes, yet suitable in vivo models remain scarce^7–11^. Since these pathogenic mutations target highly conserved functional regions, such as the RXG motif (R290H) in NUS1 and the R37/R38 residues (R37H) in DHDDS (Supplementary Figs. 11, 12)^10,21^, we hypothesized that the orthologous *C. elegans* sites, NUS-1 R223 and DHDDS R48, would be similarly critical and recapitulate the human pathogenic effects.

To test this, we used CRISPR-mediated gene editing to introduce the orthologous R223H and R48H mutations. The resulting mutant exhibited severe but distinct phenotypes, thereby confirming the physiological importance of these sites. Specifically, the NUS-1 R223H mutant was sterile, while the DHDDS R48H mutant caused larval arrest (Lva) at the L1 stage (Fig. 7a and Supplementary Fig. 13a). These severe developmental defects were consistently accompanied by significant impairment in locomotion coordination and a lipid metabolism defect (Fig. 7b and Supplementary Fig. 13b-d), all mirroring key findings in human CDG cases^63^. The more severe DHDDS R48H phenotype aligns with its predicted role in directly and fundamentally impairing the crucial *cis*-PT catalytic pocket structure^22^.

**Fig 7.**
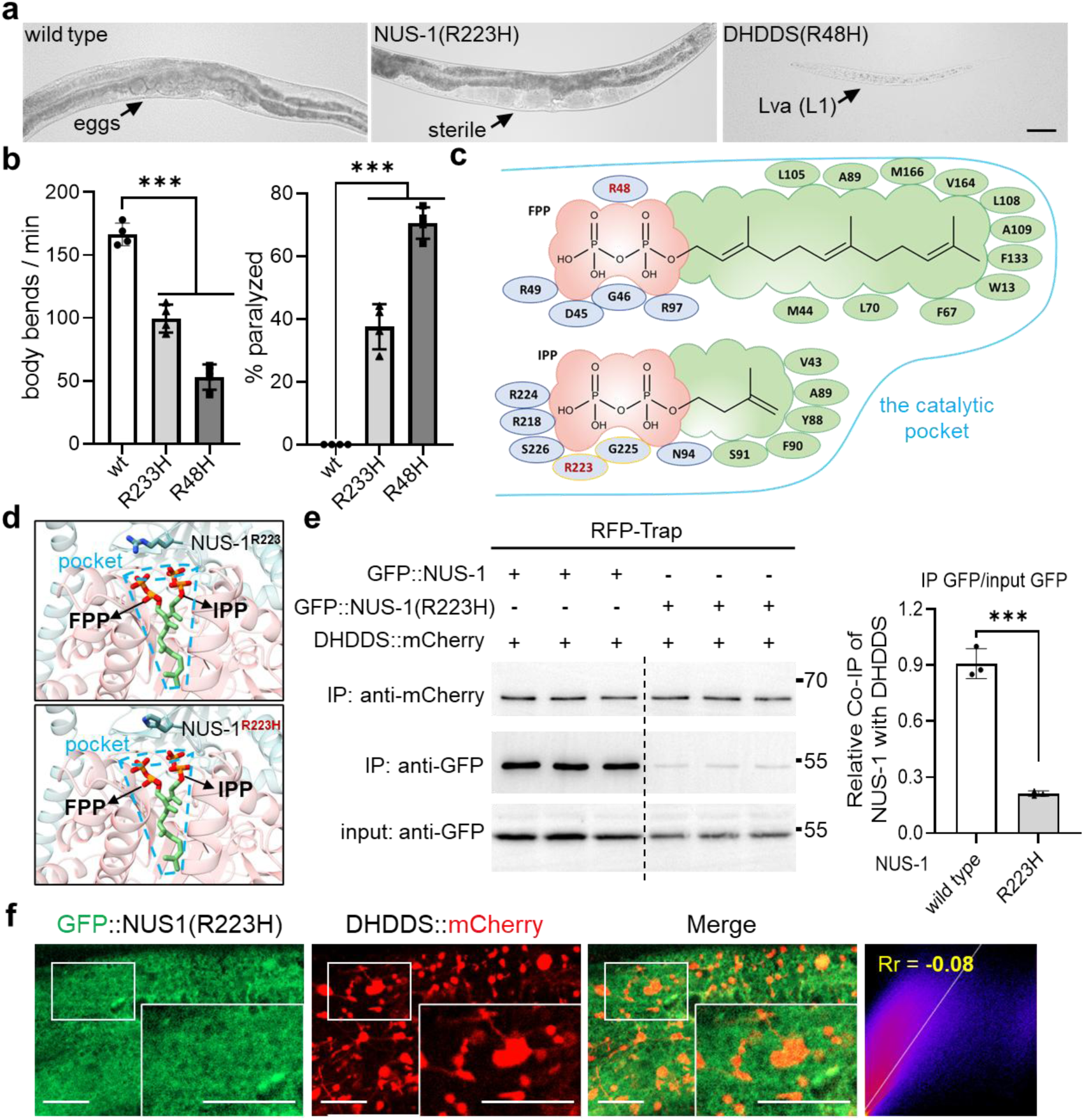
Structural and functional characterization of Human CDG-associated mutations Using *C. elegans* disease models. **a** Bright-field images illustrating the developmental defects of NUS-1(R223H) and DHDDS(R48H) mutants compared to wild-type animals. Lva, larvae arrest. Scale bar, 1 mm. **b** Impaired movement and paralysis in NUS-1(R233H) and DHDDS(R48H) CDG models at L1 stage. Movement velocity (left) measured as body bends/min (n = 15 animals/group, 4 replicates). Paralysis rate (right) defined as the percentage of animals that moved their head but not their body after a tap stimulus (n = 50 animals/group, 4 replicates). unpaired t-tests: \*\*\**p* < 0.001. **c** Predicted binding mode of FPP and IPP in the *C. elegans cis*-PT catalytic pocket. The hydrophobic pocket is outlined in blue. Substrates feature FPP (top) and IPP (bottom): negatively charged phosphate groups are highlighted in red, and the carbon backbone in green. Positively charged residues stabilizing the phosphates are shown in blue background, specifically NUS-1 (yellow circle) and DHDDS (blue circle). Hydrophobic interactions with the backbone are mediated by DHDDS residues (green background). Pathogenic orthologous residues R48 (DHDDS) and R223 (NUS-1) are highlighted in red font. **d** Structural comparison of the *cis*-PT catalytic pocket in NUS-1 wild-type (top) and the R223H mutant (bottom). The NUS-1 subunit is cyan and DHDDS is salmon (transparent cartoon view). The catalytic pocket, indicated by a dotted blue line, contains FPP (left) and the IPP (right) substrates. The key residue R223 is highlighted in stick representation, showing its position above the pocket and the predicted structural impact of the R223H substitution. **e** Representative Western blot showing and quantification from a co-Immunoprecipitation (Co-IP) assay showing the interaction between DHDDS::mCherry and GFP::NUS-1 or GFP::NUS-1(R223H). Lysates from animals expressing these fluorescently tagged proteins were subjected to IP using RFP-TRAP, followed by Western blotting with antibodies against GFP and mCherry (n = 3 per group, unpaired t-tests: \*\*\**p* < 0.001). **f** Confocal image showing colocalization of DHDDS::mCherry with GFP::NUS-1(R223H). Rr < 0, negative colocalization. Scale bar, 10 μm.

We utilized AutoDock4 software and structural models to predict the substrate-binding mechanisms^64,65^. The prediction confirmed that FPP and IPP bind conservatively within the DHDDS-formed catalytic pocket (Fig. 7c)^22^. Specifically, the negatively charged phosphate groups of FPP form three key electrostatic interactions with the R48 side chain within the pocket (Supplementary Fig. 13e). The R48H mutation was predicted to disrupt these stabilizing interactions, severely impairing pyrophosphate headgroup stabilization. Conversely, NUS-1 R223 acts as a structural lid above the pocket without direct substrate interaction (Fig. 7d). The R223H substitution was predicted to prevent the substituted Histidine residue from properly ’covering’ the pocket opening.

We then validated these structural predictions using co-IP. The NUS-1 R223H mutation severely compromised the physical interaction between NUS-1 and DHDDS (Fig. 7e and Supplementary Fig. 13f), while the DHDDS R48H mutation did not (Supplementary Fig. 13g). This indicates that the NUS-1 R223H substitution impacts catalytic pocket architecture, resulting in reduced NUS-1/DHDDS interaction affinity and the sterile phenotype.

Importantly, a cycloheximide chase assay confirmed that neither mutation affected protein turnover, demonstrating that the loss of function is not due to protein instability (Supplementary Fig. 13h, i)^66^. Crucially, the impaired interaction caused by NUS-1 R223H also simultaneously abolished its colocalization with DHDDS (Fig. 7f and Supplementary Fig. 13j). As the soluble DHDDS subunit is recruited to the ER membrane by anchored NUS-1 to form the *cis*-PT complex, this provides a clear mechanism for the loss of function. These results unequivocally demonstrate two independent mechanisms for *cis*-PT dysfunction: interface disruption (NUS-1 R223H) and catalytic impairment (DHDDS R48H). This work collectively establishes a clear mechanistic distinction and provides a robust *C. elegans* platform for investigating the molecular etiology of human CDG-associated disorders.

## Discussion

Our study utilizes the genetic tractability of *C. elegans* to identify the conserved NUS-1/DHDDS complex as a critical dual regulator linking N-glycosylation to global lipid and lysosomal homeostasis. By overcoming the limitations of embryonic lethality associated with mammalian germline knockouts, we provide a whole-organism analysis that reveals an essential crosstalk between N-glycosylation, lipid metabolism, and lysosomal integrity. This work establishes a valuable in vivo platform for dissecting the multi-faceted pathophysiology of CDGs (Fig. 8).

**Fig 8.**
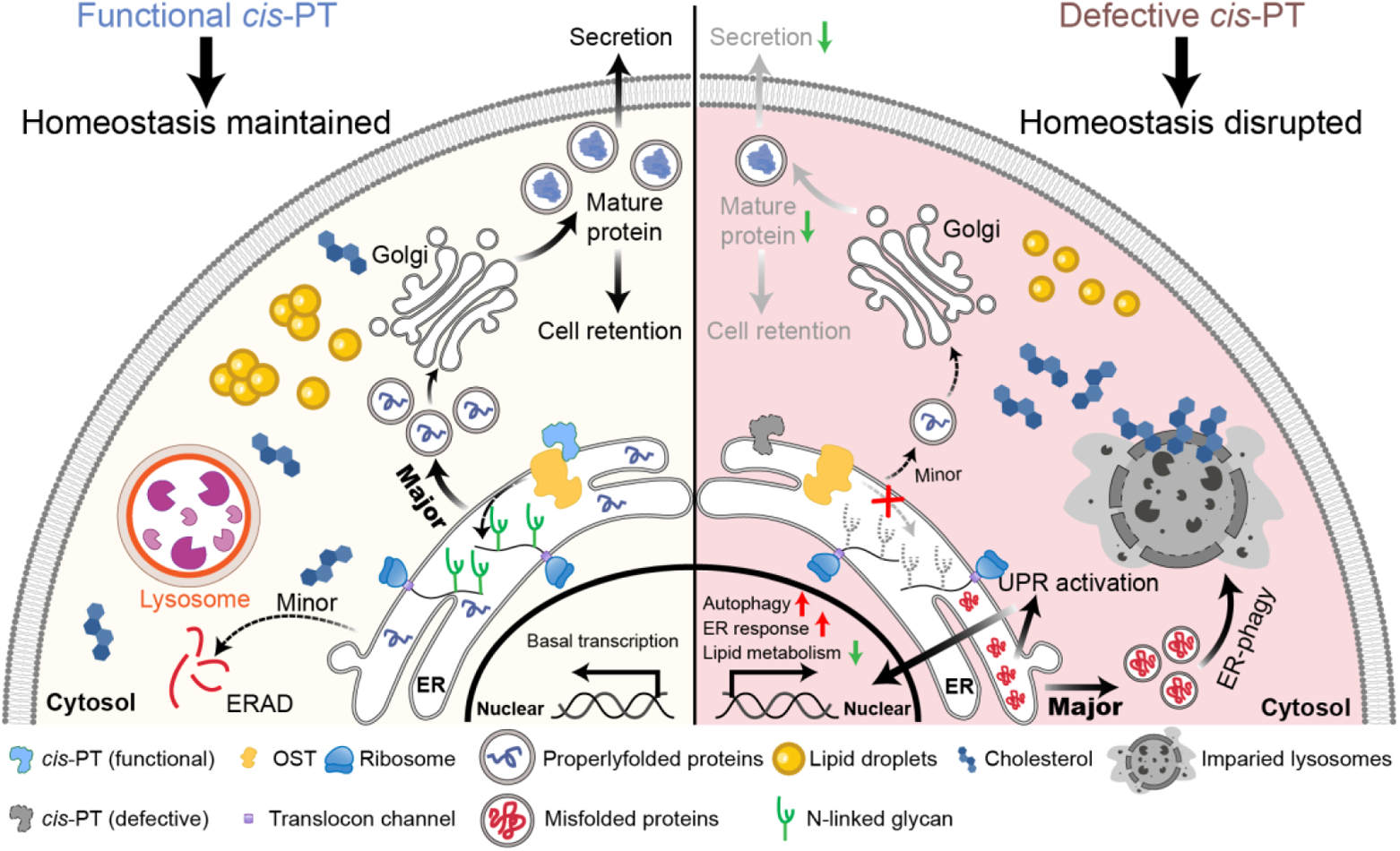
Schematic diagram of defective *cis*-PT influencing cellular homeostasis. In wild-type organisms, the functional *cis*-PT complex is essential for producing the correct polyprenyl phosphate substrate required for N-glycosylation by the OST complex in the ER. This ensures efficient processing and routing of properly folded proteins for secretion or retention, with minor misfolded proteins cleared via ER-associated protein degradation (ERAD). When the NUS-1/DHDDS complex is defective, the resulting N-glycosylation defect and impaired substrate availability leads to the accumulation of misfolded proteins in the ER lumen. This proteotoxic stress activates the unfolded protein response (UPR^ER^ ), intensifying compensatory degradation pathways (ERAD and ER-phagy), and disrupting protein secretion and retention. This systemic breakdown of proteostasis is further exacerbated by concurrent impairment of lysosomal homeostasis and lipid metabolism pathway, culminating in cholesterol accumulation within cells. Collectively, these cascading defects disrupt the overall homeostasis of the organism.

Despite limited sequence identity, our findings underscore the functional conservation of the *cis*-PT complex across evolutionary timescales. While human NUS1 possesses three N-terminal TM domains^10^, its *C. elegans* ortholog utilizes a single, non-cleavable TM segment for ER anchoring. More significantly, our data illustrate an evolutionary refinement of the OST machinery. Unlike vertebrates, which evolved specialized OST-A (STT3A-containing, co-translational) and OST-B (STT3B-containing, post-translational) complexes to accommodate a complex secretome^41,67^, *C. elegans* maintains a streamlined system centered on a single STT-3 catalytic subunit and its auxiliary factor ZK686.3 (the common ancestor of human MAGT1 and TUSC3)^19^. This architectural parsimony in *C. elegans* facilitates both nascent glycan transfer and stress-responsive glycosylation, l reflecting metabolic adaptation to a shorter lifespan and reduced protein flux compared to higher eukaryotes.

A central discovery of this work is the direct physical association between the *cis*-PT complex and the core OST machinery. This interaction, primarily mediated by NUS-1 and the STT-3/RIBO-1 subunits, suggests a mechanism of proximal substrate channeling. In this model, the *cis*-PT complex functions as an integrated biosynthetic module that directly supplies newly synthesized Dol-P to the OST assembly line. This spatial coordination likely bypasses the kinetic bottlenecks of lipid diffusion within the ER membrane, thereby ensuring maximal glycosylation efficiency. Our structural modeling further predicts that this interface is highly conserved in humans, involving hNUS1 and the OST-B specific subunits STT3B and MAGT1. Notably, cross-referencing our structural model with UniProt clinical variants reveals that several interface residues correspond to human pathogenic sites (NUS1 S146Y, DHDDS E307Q, and MAGT1 T251M). This suggests that certain CDG phenotypes may arise not from intrinsic enzymatic deficiency, but from the functional uncoupling of precursor production and glycan assembly.

This primary glycosylation defect extends to a catastrophic breakdown in cellular proteostasis and lipid homeostasis. We demonstrate that the instability of critical glycoproteins, such as lysosomal membrane protein LMP-1 and various collagens, leads to lysosomal membrane permeabilization and diminished degradative capacity. This lysosomal collapse triggers a robust, yet ineffective, compensatory activation of autophagy. Crucially, the observed lipopenia arises from the impaired nuclear translocation of the master lipogenic transcription factor SBP-1. This defect is likely driven by the under-glycosylation of its essential escort protein, SCAP (SCP-1)^68^. Such a regulatory link provides a molecular basis for the systemic metabolic disturbances observed in CDG patients, where clinical manifestations often extend far beyond primary glycosylation deficits^69^.

Finally, our *C. elegans* CDG models, which harbor the NUS-1 R223H and DHDDS R48H mutations, recapitulate the developmental and locomotor defects observed clinically. We show that these mutations disrupt the *cis*-PT complex through distinct molecular mechanisms: interface destabilization (R223H) and catalytic pocket impairment (R48H). This mechanistic heterogeneity underscores the necessity for precision medicine approaches in treating CDGs. Future investigations utilizing mammalian systems, such as proximity labeling or cryo-electron microscopy, will be vital to confirm the conservation of this supercomplex and to determine whether stabilizing the *cis*-PT/OST can restore glycosylation flux in disease contexts. Collectively, these findings fill a critical gap in our understanding of N-glycosylation substrate transport and establish *C. elegans* as a powerful system for pharmacological screening and the study of glycoprotein-related disorders.

## Methods

### *C. elegans* strains and maintenance

All *C. elegans* strains, except those subjected to RNAi analysis, were maintained at 20°C on standard nematode growth medium (NGM) plates seeded with *E. coli* OP50. The N2 Bristol strain served as the wild-type control. Transgenic lines were generated by microinjection of Day 1 adult gonads with co-microinjected plasmids (Supplementary Table 2). All resulting transgenic worms were consistently maintained and screened at 20°C, except for mD4H transgenic lines, which required screening and subsequent RNAi observation experiments to be performed at 25°C^70^. CRISPR/Cas9 edited strains were generated by SunyBiotech and confirmed by Sanger sequencing (Supplementary Table 2). Integrated alleles *bmsIs8* and *bmsIs117* were generated via UV radiation (Supplementary Table 1).

### RNAi treatment

RNAi bacteria were cultured in LB medium containing 100 μg/mL ampicillin at 37°C for 14 hours. The bacteria were then concentrated 5-fold by centrifugation and resuspended before being inoculated onto NGM plates containing 50 μg/mL ampicillin and 1 mM IPTG. After induction at room temperature for 24 hours to ensure sufficient dsRNA production, worms were transferred onto the RNAi plate. For L4 progeny analysis, animals at L4 stage were placed on RNAi plates and their F1 progeny at L4 stage were analyzed 4 days later. For P0 adult analysis, eggs were placed directly onto RNAi plates, and analysis were performed on the resulting adults 5 to 7 days later.

### Plasmid construction

All overexpression plasmids were constructed using the pPD95.75 backbone. The 1400 bp *nus-1* promoter was used to drive the expression of either the full-length sequence for *nus-1* genomic DNA or a truncated version (Δ2-33aa), each fused with a 3xFLAG tag. Transgenic strains were generated by microinjection of the plasmids, and the resulting protein expression was confirmed via Western blot analysis.

### Microscopy and phenotypic imaging

All confocal images were acquired using a high-resolution laser scanning confocal microscope (Carl Zeiss, LSM980). All bright-field and non-confocal fluorescence images were captured using an upright fluorescence microscope (Nexcope, NE910). For live fluorescence imaging, worms from different treatment groups but at the same developmental stage (L4 or adult) were randomly selected and immobilized on 2.5% agarose pads. Immobilization was achieved by placing in 10 mM levamisole (dissolved in M9), which was placed onto the pad prior to imaging. For gross phenotypic imaging, animals were selected without balancer (L4 or Day 1 adult stage for viable mutants). A minimum of five worms per strain were observed, and representative images were selected for display using a fluorescence stereo microscope (Nexcope, NSZ818).

### Quantification of polyglutamine aggregates

Polyglutamine aggregate puncta were quantified in both Q35::YFP and Q40::YFP transgenic strains. L4 or Day 6 adult worms from all treatment groups were randomly selected and observed while moving on NGM plates using a fluorescence stereo microscope (Nexcope, NSZ818). The number of YFP-labeled puncta were manually counted along the entire body length of each worm, and the resulting data were subjected to statistical analysis.

### Structural prediction and visualization

Protein sequences for *C. elegans* NUS-1 (NP_001379086.1) and DHDDS (NP_001023351) were utilized for structural modeling. Substrate-binding conformations for FPP and IPP were predicted using AutoDock v.4.2.6. All structural visualizations and molecular edits were performed using ChimeraX v.1.10.1. All protein complex structures in the absence of substrates were predicted using AlphaFold3. For *C. elegans* OST structural analysis, the following amino acid sequences were retrieved from NCBI: STT-3 (NP_498362.1), RIBO-1 (NP_501065.2), OSTD-1 (NP_491624.1), OSTB-1 (NP_495655.1), DAD-1 (NP_491889.1), ZK686.3 (NP_498691.1), OSTF-4 (NP_001367095.1), and TMEM-258 (NP_504794.1). For the human OST complex, structural data were obtained from the Protein Data Bank (PDB: 6S7T). Structural comparisons between the *C. elegans* and human *cis*-PT were conducted by importing both models into ChimeraX and performing a best-match alignment to determine structural homology. The *cis*-PT/OST supercomplex structures for both species were predicted using AlphaFold3, with final models processed and visualized in ChimeraX.

### Measurement of *C. elegans* body length and width

Body length and width were quantified using animals at L4 stage. Body length was measured from the anterior tip of the mouth to the tail region posterior to the anus. Body width was defined as the distance perpendicular to the body axis, measured specifically at the vulva.

### Quantification of locomotion and paralysis

The locomotion activity was quantified by measuring body bends per minute. ≥ 15 worms at the same stage, cleaned of OP50 bacteria, were collected and placed on the 2.5% agarose pad with a 1 μL M9 drop. Locomotion was recorded for 1 min under a bright-field microscope. A single body bend was defined as a maximal deflection of the region posterior to the pharynx towards the opposite direction from the previous deflection. The percentage of paralyzed animals (% paralyzed) was determined by gently prodding animals on OP50-seeded NGM plates with a platinum worm picker. Paralysis was scored if the worm, upon being stimulated near the head/anterior region, failed to initiate body movement.

### NUC-1::pHTomato reporter analysis

The *C. elegans* strain expressing the P*_hs_*NUC-1::pHTomato reporter was used to assess lysosomal acidification. P0 animals were subjected to RNAi treatment, and the resulting F1 progeny at L4 stage were first subjected to a 32°C heat shock for 1 h to induce expression. Following induction, animals were allowed to recover at 20°C for 4 hours before imaging^71^. The average intensity of pHTomato was then quantified per puncta in the hypodermis using ImageJ software. A minimum of 360 puncta were analyzed for each strain.

### Co-immunoprecipitation (Co-IP) and mass spectrometry

Worms were collected from NGM plates, washed with M9 buffer, and homogenized using a glass pestle in 1 mL IP buffer (Beyotime, P0013) supplemented with 1 mM PMSF and 1x Protease Inhibitor Cocktail (Bimake, B15001). Homogenates were incubated for 2 h at 4°C with rotation to ensure complete lysis. Lysates were centrifuged at 5,000 rpm for 1 min at 4°C. The resulting supernatant was pre-cleared by incubating with Binding Control Agarose (Chromotek, bmab-20) for 20 min at 4°C. The pre-cleared supernatant was then incubated with GFP-Trap Magnetic Agarose (Chromotek, gtma-20) or RFP-Trap Magnetic Agarose (Chromotek, rtma-20) for 3 h at 4°C. Beads were washed three times with 1x PBS and eluted by boiling in 1x SDS sample buffer. Input samples were prepared with 80 μL of the pre-cleaned supernatant with 20 μL of 5x SDS sample buffer and boiling. For mass spectrometry analysis, eluted co-IP samples were separated by SDS-PAGE. The gel was stained using a commercial silver staining kit (Beyotime, P0017S), and specific protein bands unique to the experimental group were excised and submitted for mass spectrometry analysis.

### Cycloheximide (CHX) pulse-chase assay

For protein stability analysis, HEK293T cells were transfected with plasmids expressing NUS-1 or DHDDS, tagged with a GFP or 1xFLAG, respectively. Forty-eight hours after transfection, cells were treated with 50 µg/mL CHX for the indicated time points. Following CHX treatment, cells were harvested and protein levels were analyzed by Western blotting.

### Western blotting

For Western blot analysis, proteins were separated by SDS-PAGE (using 10% or 12% resolving gels, depending on the size of target protein) and subsequently transferred to PVDF membranes. Membranes were blocked with 5% non-fat milk in 1x TBS containing 0.1% Tween-20 and incubated overnight at 4°C with primary antibodies. Membranes were then incubated with secondary antibodies at room temperature before visualization using SuperPico ECL Chemiluminescence Kit (Vazyme, E422) on the Imaging System (Tanon, 5200). The following primary antibodies were used: anti-GFP (Abbkine, ABT2020; abclonal, AE011), anti-mCherry (Invitrogen, M11217), anti-Histone H3 (abcam, ab1791), and anti-tubulin (Sigma, T5168).

### PNGase F deglycosylation assay

Worms were lysed using a commercial lysis buffer (Beyotime, P0013), and protein concentration was determined by BCA assay (Beyotime, P0010). 40 μg of total protein was incubated with PNGase F (NEB, P0704S) for 4 h at 37°C. The control group, containing all reagents except PNGase F, was incubated simultaneously under the same conditions. All samples were subsequently analyzed by Western blotting.

### Quantification of PVD dendrite morphology

PVD dendrite morphology was quantified by confocal microscopy (Carl Zeiss, LSM980). Animals from all groups at L4 stage were aligned in the same orientation on agarose pads. The status of the first ten secondary dendrites emanating from the PVD soma were recorded. A secondary dendrite was classified as “abnormal” if it exhibited any of the following morphological defects: the presence of punctate neuronal knots/neuropils along the dendrite, failure to properly generate tertiary dendrites, or inappropriate extension and fusion with an adjacent secondary dendrite.

### Oil Red O staining and quantification

Neutral lipid levels were assessed using Oil Red O (ORO) staining (Sigma, O1391). Worms were collected, washed in 1x PBS, and fixed for 1 h using 4% paraformaldehyde dissolved in 2x MRWB buffer^55,72^. To dehydrate the tissue, samples were incubated in 60% isopropanol for 10 min at room temperature. For staining, ORO stock solution (saturated isopropanol solution) was diluted to 60% (v/v) with water, mixed for 1 h, and filtered through a 0.45 μm organic membrane. Samples were stained with the filtered 60% ORO solution for 1.5 h. After washing in PBST and PBS, bright-field images were acquired, and the final quantification of neutral lipids was performed by measuring the average optical density (AOD) of the stained area using ImageJ.

### Filipin staining for unesterified cholesterol

Unesterified cholesterol was visualized using Filipin (Sigma, SAE0087), following previously established protocols^70,73^. Worms were collected and washed in ice-cold double-distilled water (ddH_2_O). Samples were fixed by adding 1 mL of pH 4.5 ethanol solution and subjected to 3 times freeze-thaw cycles using liquid nitrogen and ethanol. Following the final thaw, the fixative was washed twice with ddH_2_O. Quenching was performed by incubating samples in 1 mL of 100 mM NH_4_Cl solution for 30 minutes at room temperature, followed by overnight incubation at 4°C. The NH_4_Cl solution was washed twice with ice-cold ddH_2_O. The pellet was resuspended in 30 μL of residual liquid, and a 30 μL of Filipin working solution (final concentration 0.5 mg/mL) was added. Samples were stained for 2 hours at room temperature, protected from light. After two final washes with ice-cold ddH_2_O, the precipitate was mounted on 2.5% agarose pads. Imaging was conducted using the DAPI channel, and slides were mounted with a fluorescence mounting medium (Invitrogen, P36982) for preservation.

### Quantitative real-time PCR (qRT-PCR)

Total RNA was extracted from approximately 200 L4 stage animals collected in 500 μL Trizol (Invitrogen, 15596026CN). Samples were subjected to three cycles of freeze-thaw using liquid nitrogen and a 65°C metal bath, followed by standard isopropanol precipitation. 1200 μg of total RNA from each sample was reverse-transcribed using a commercial kit (Vazyme, R312-01). qRT-PCR was performed using ChamQ SYBR Color qPCR Master Mix (Vazyme, Q411-02). All gene expression changes were normalized to the expression of the housekeeping gene *act-1*.

### Dual lysosomal staining

Lysosomal morphology and acidity were assessed using LysoSensor Green DND-189 (LSG, Yeasen, 40767ES50) and LysoTracker Red DND-99 (LTR, Yeasen, 40739ES50) staining. Worms were collected and washed in room temperature M9 buffer. 100 μL of concentrated worm suspension was incubated with 1 μL of the stain mixture (final concentration 10 μM for both dyes). Staining was carried out for 1 h at 20°C in the dark. Following staining, worms were recovered for 2 h at 20°C on NGM plates seeded with OP50, protected from light. Recovered worms were then mounted on 2.5 % agarose pads for confocal observation. The relative fluorescence intensity ratio of LSG to LTR was quantified using ImageJ software.

### Statistical analysis and reproducibility

All data were analyzed using GraphPad Prism 10.4.1. Unless otherwise noted, data are presented as the mean ± standard deviation (S.D.). Statistical significance (P values) was calculated using an unpaired two-sided t-test for comparisons between two groups. Statistical significance was defined as *p* < 0.05 and indicated by a single asterisk (*). Double asterisks (**) denoted *p* < 0.01, and triple asterisks (***) indicated *p* < 0.001. All experimental results were independently repeated a minimum of two times with similar outcomes. Only representative images are displayed in the figures.

### Data availability

All data supporting the findings of this study are available within the accompanying Source Data file provided with this paper, including data presented in the main manuscript and the supplementary Information. All unique reagents and strains generated in this study are available from the corresponding author upon reasonable request and the completion of a standard Material Transfer Agreement (MTA).

## Supporting information

Supplementary Table 1. C. elegans strains used in this study.

Supplementary Table 2. Primers used in this study

## Acknowledgements

We thank the Caenorhabditis Genetics Center (CGC), National Bioresource Project (NBRP), the Mitani lab, Dr. Xiaochen Wang (Southern University of Science and Technology), Dr. Wei Zou and Dr. Caiyong Chen (Zhejiang University), Dr. Ye Tian and Dr. Yidong Shen (Chinese Academy of Sciences), Dr. Dengke Ma (UCSF), and Dr. Bin Liang (Yunnan University) for providing strains. We thank Dr. Dandan Wang (Fudan University) for providing plasmids, and Dr. Qinghua Zhou (Jinan University) for technical assistance with Oil Red O staining. We are especially grateful to Dr. Yile Zhai, Dr. Tiantian Wang, and Dr. Mengxiao Wu (Shandong University) for their critical reading, constructive feedback, and valuable suggestions on the manuscript.

## Funding

This study was supported by grants from the National Natural Science Foundation of China (32170781 to Z.Z.), the Natural Science Foundation of Shandong Province (2023HWYQ-014, tsqn202306056, ZR2021QC023, and 2021JK032 to Z.Z.).

## Author contributions

Conceptualization: M.G, C.L, W.L, P.Z, Z.Z; Data Curation: M.G, W.Z, W.L, P.Z, Z.Z; Formal Analysis: M.G, W.L, P.Z, Z.Z; Funding Acquisition: Z.Z; Investigation: M.G, Y.D, Y.Z, C.J, M.W, T.W, W.Z, W.L, P.Z, Z.Z; Methodology: M.G, Y.Z, C.J, W.Z, W.L, P.Z, Z.Z; Project Administration: C.L, W.L, P.Z, Z.Z; Resources: M.G, W.L, P.Z, Z.Z; Supervision: C.L, W.L, P.Z, Z.Z; Validation: M.G, W.L, P.Z, Z.Z; Visualization: M.G, W.Z, W.L, P.Z, Z.Z; Writing – Original Draft: M.G, Z.Z; Writing – Review & Editing: M.G, W.L, P.Z, Z.Z.

## Competing interests

The authors declare no competing interests.

## Notes

### Competing Interest Statement

The authors have declared no competing interest.

